# Characterization of strawberry (*Fragaria vesca*) sequence genome

**DOI:** 10.1101/244418

**Authors:** Ao Li, Lide Chen, Zhongjie Liu, Mengjie Cui, Lingfei Shangguan, Haifeng Jia, Jinggui Fang

**Affiliations:** Nanjing Agricultural University, Nanjing, 210095, China.

**Keywords:** strawberry, exon, intron, Structural features, Correlation analysis, gene expression

## Abstract

**Background:** In order to understand strawberry genes’ structure and evolution in this era of genomics, it is important to know the general statistical characteristics of the gene, intron and exon structures of strawberry and the expression of genes on different parts of strawberry genome. In the present study, about 32,422 genes on strawberry chromosomes were evaluated, and a number of bioinformatic softwares were used to analyze the characteristics of genes, exons and introns, expression of genes in different regions on the chromosomes. Also, the positions of strawberry centromeres were predicted.

**Results:** Our results showed that, there are differences in the various features of different chromosomes and also vary in different parts of the same chromosome. The longer the number of genes, the longer the length of chromosome. The average length of genes is about 2809bp and the length of the individual gene is 0–2000bp with 5.3 exons and 4.3 introns per gene. The average length of the exon was 229bp and the intron was 413bp. Among the evaluated genes, ehe intronless gene accounted for 20.05%. Consistently a same trend with the expression levels of the same parts of the gene on a chromosome in different organizations was observed. Finally, the number of genes was positively correlated with the number of intronless, and there was a negative correlation of the length of the gene. The length of the gene depends primarily on the length of the intron, and the length of the exon has little effect on it. The number of exons was negatively correlated with the length of the exons, and the intron was also true.

**Conclusion:** The results of this investigation could definitely provide a significant foundation for further research on function analysis of gene family in Strawberry.

## Background

Since the dawn of the genomic age accurate prediction of protein coding (and other) genes has been a central problem of biology in genomics [1–3]. This field has advanced quickly in a short time, which largely due to the surge in whole genome sequencing, the development of bioinformatics tools and the associated wealth of new data, over the period. However, accurate genome annotation is an extremely difficult problem which requires balancing of false negatives and positives, and accuracy versus time constraints [4]. Strawberry (*Fragaria vesca*) is an economically important fruit crop in the world, and its whole genome sequence was reported in 2011 [5] which make possible for the characterization of strawberry genes and accurate genome annotation. In this investigation, we examined the distributions of genes, exons, introns and expression of the genes on the 7 strawberry chromosomes and discern the correlations between them which are most fundamental for a quantitative view of strawberry genome structure. These findings give a brief view of some strawberry genome characters and help improve gene structure prediction by computational methods by providing better understanding of factors that govern genome design and architecture [6].

The Characterization of many plant genomes have been focused on during the past years. Studies on gene density about the known rice contains more than twice as many genes as those of *Arabidopsis thaliana* was reported [7]. In addition, we know that the genes of higher eukaryotes are mostly broken, and the lengths of genes between different species are very different. For example, between *E. coli* and humans the average gene content meets by nearly two orders of magnitude [8]. The length of the genes in same species are also different, such as the length of the human protein-coding genes which were widely distributed and range from a few hundred bases up to a few million [9]. Previous studies of gene lengths could predict human disease [10] and also had a certain effect on gene expression [11]. The analysis of gene length has kept being focused on the gene structure and evolution. The intron-exon structures of genes were also an interesting character that had been studied in genomics field. Complete statistical description of intron-exon structures is imperative for the theoretical study of the origin and evolution of genes and genomes. An exon of any segment of an interrupted gene is represented in the mature RNA product, and an intron being the part of eukaryotic genes without translation [12]. However, prokaryotic genes lack introns although intronless genes are a characteristic feature of prokaryotes, eukaryotes have both intron-containing and intronless genes. The percentage of intronless genes varies from 2.7% to 97.7% of the genes in eukaryotic genomes [13]. Many authors have published the analysis of some characteristics of nuclear introns in various organisms [14–16].

The expression levels of genes can indicate the role of the genes studied, and the gene expression levels are also distributed with a wide range across the genes of the organism and related with the gene structures. For example a strong correspondence was reported between the level of a genes expression and the length of its introns [17]. Highly expressed genes tend to have shorter introns, likely reflecting their adaptation for high transcription efficiency; gene length was examined in a set of constitutively expressed genes [18]. Various investigators found that genes with the highest level of expression tend to be those that were also the shortest in length. Grishkevich et al. demonstrated that a requirement for high expression imposes a considerable constraint on gene length [9].

With the rapid development of genetic informatics and the completion of genome sequencing of wild strawberry [5]. Recent advances have made available an enormous resource of data for the study of strawberry genomes, and strawberry molecular biology has become more and more popular. Strawberry represents an emerging system for the study of gene and genome evolution and processes related to plant physiology, development and crop production, which benefited from its small stature, rapid seed-to-seed cycle, transformability and miniscule basic genome. Therefore, a study on the number and length of the gene, introns and exons of strawberry genes, the distribution of intronless genes, gene expression and correlation among them can be important for deeper study of strawberry genomics.

## Methods

### Data sources

The strawberry data used in this study were downloaded from the website of Genome Database For Rosaceace (https://www.rosaceae.org/organism/Fragaria/vesca). The data of strawberry electronic expression analysis to predict the position of strawberry chromosome centromere were obtained from the website of Strawberry Genomic Resources (http://bioinformatics.tows-on.edu/strawberry/newpage/Search_By_Gene_No.aspx).

### Analysis tools

UltraEdit software was used for data selection of the genes, exons and introns and processing. Echarts and Origin 7.0 software were used to draw the chart. Rstudio software was employed for heatmaps analysis with RStudio version using Heatmaps.2 from the ‘gplots’ package. The specific execution of the R script and the role of the command line is as follows:

~~~
Library (ggplot2) and Library (reshape2)

mat <-read.table (“.txt”,header=TRUE, sep=“”)

mat

m = melt (mat) m

g = ggplot (m, aes (x=variable, y=ID, fill=value)) +xlab (‘lg (Expression Value)’) +ylab (“Gene ID”)

g1=g+geom_tile (); print (g1)

g2=g1+geom_tile (color=“white”, size=0.1); print (g2)

g3=g2+scale_fill_gradient (low=‘green’, high=‘red’);print (g3)

g4=g3+geom_text (aes(label=round(value,2)), angle=45);print (g4)

#400*400
~~~

### Sequence data chosen and processed

Five segments of 2M length were chosen from each strawberry chromosome: upper, middle upper, middle, middle lower and lower (Fig. 1). Totally, 35 segments were analyzed further to characterize the strawberry genome, which can be used in comparison between different chromosomes and segments on the same chromosome. The total length of the five segments chosen on each chromosome covered 43.1%, 32.6%, 30.2%, 30.9%, 34.2%, 25.8% and 42.4% of their corresponding chromosomes, respectively. Totally, 33.4% of the strawberry genome was studied. Then through the R software carried out heatmap analysis in order to compare the average expression of the genes location on different chromosome regions in different tissues. Finally the partial correlation coefficientbetween various indicators was computed using the SPSS function.

**Figure 1.**
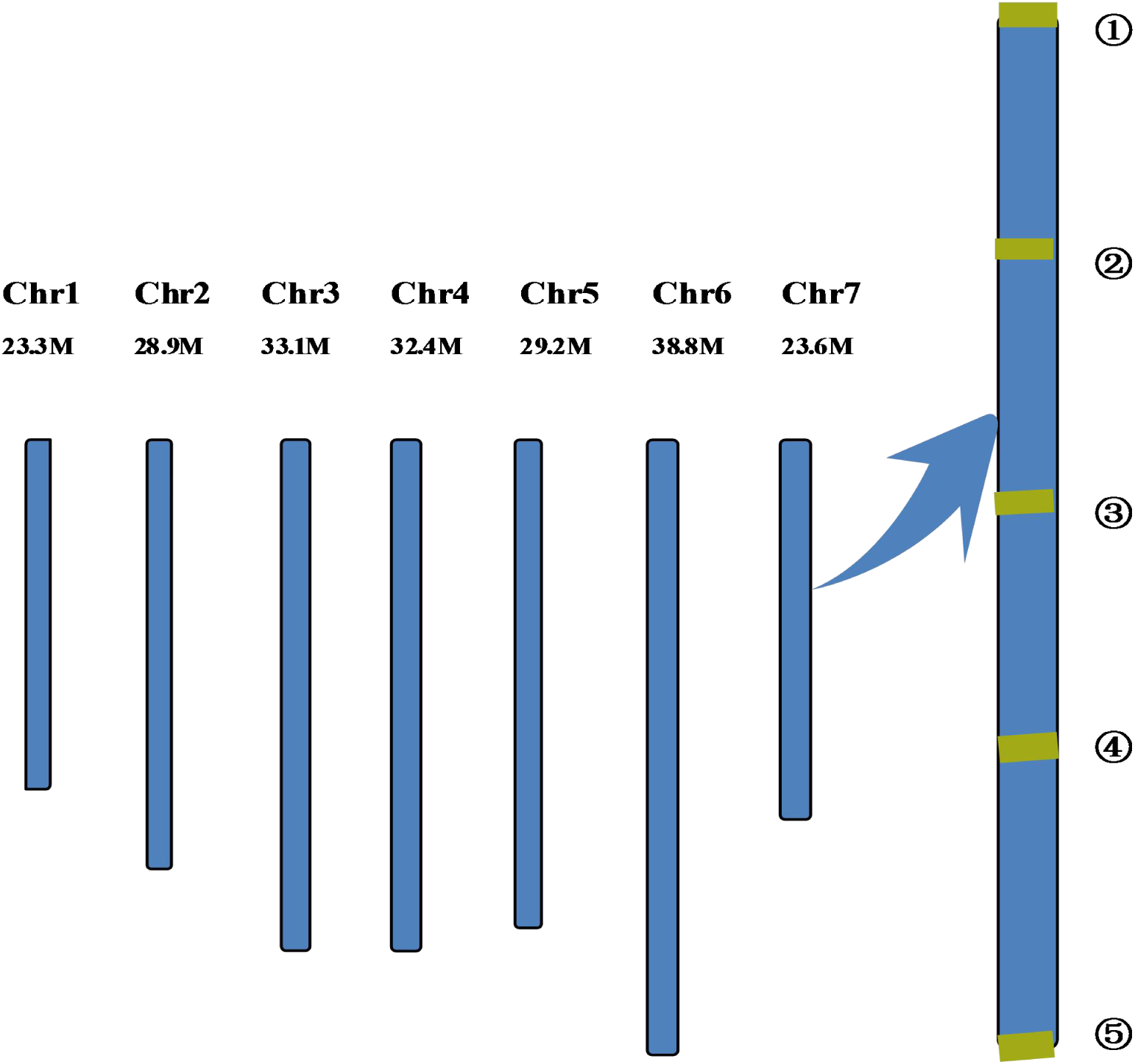
Location of genes in different chromosomal regions Note: ①, ②, ③, ④,⑤: upper, middle upper, middle, middle lower and lower 23.3M, 28.9M, 33.1M, 32.4M, 29.2M, 38.8M, 23.6M represents the length of each chromosome

## Results

### Distributions of genes, exons and introns in strawberry genome

The contents of the characteristics of the gene, exon and intron number and length which were analyzed in this study is summarized (Table 1). A total of 11,574 host genes were selected, among which the average number of the genes on all the 35 regions was 331 with a range from 195 to 503. The region with lowest gene number is the 1–5 regions on Chr1 with 195 annotated genes, and the highest region is the 4–5 on Chr4 with 503 genes. The average length of all the selected genes was about 2,809 bp, and the length variation of the genes on the different regions was similar. The longest annotated gene is 30,412 bp, a gene regulating, and the shortest gene was 92 bp in size. The average number of exons in a gene was about 4–6 and the mean number was 5.3 exons per gene. The average exon length was about 229 bp. Conversely, the average intron size is about 413 bp.

**Table 1.** Gene, exon and intron distributions for strawberry genome

| Chromosome | fragment | genes number | Exon number<br>(Avg number ) | Intron number<br>(Avg number) | Gene length(bp)<br>(Avg length ) | Exon length(bp)<br>Avg length | Intron length(bp)<br>Avg length |
| --- | --- | --- | --- | --- | --- | --- | --- |
| Chr1 | 1-1 | 426 | 1-25 (5.3) | 0-24 (4.3) | 259-23063(2695.3) | 5-2568(279.3) | 24-7549(339.8) |
|  | 1-2 | 395 | 1-19 (5.0) | 0-18 (4.0) | 232-19136(2860.6) | 27-2408(268.7) | 28-3427(294.0) |
|  | 1-3 | 328 | 1-23 (4.5) | 0-22 (3.5) | 92-12056 (2468.4) | 21-2201(258.0) | 73-3603(425.6) |
|  | 1-4 | 290 | 1-28 (4.5) | 0-27 (3.5) | 214-12609(2689.7) | 5-2066(244.6) | 31-7520(407.6) |
|  | 1-5 | 198 | 1-15 (4.8) | 0-14 (3.8) | 191-21180(3280.2) | 19-3005(217.3) | 29-8394(624.7) |
| Chr2 | 2-1 | 289 | 1-58 (6.4) | 0-57 (5.4) | 227-19001(2745.5) | 14-1895(202.3) | 35-7803(440.8) |
|  | 2-2 | 260 | 1-14 (4.2) | 0-13 (3.2) | 237-18140(2555.9) | 6-1713(225.6) | 55-9631(578.1) |
|  | 2-3 | 343 | 1-17 (4.1) | 0-16 (3.1) | 270-14608(2690.4) | 7-1824(265.7) | 42-7711(466.6) |
|  | 2-4 | 371 | 1-31 (6.2) | 0-30 (5.2) | 190-15411(2847.5) | 23-2669(231.4) | 30-6068(364.5) |
|  | 2-5 | 431 | 1-31 (6.2) | 0-30 (5.2) | 270-14608(2725.9) | 5-2654(204.3) | 29-4402(316.5) |
| Chr3 | 3-1 | 415 | 1-23 (6.0) | 0-22 (5.0) | 259-24149(2635.6) | 19-2087(236.3) | 40-5701(321.0) |
|  | 3-2 | 315 | 1-25 (6.0) | 0-24 (5.0) | 204-19542(2937.7) | 9-2966(195.7) | 36-3931(401.1) |
|  | 3-3 | 231 | 1-49 (6.4) | 0-48 (5.4) | 249-21309(2898.3) | 6-3458(213.4) | 40-9757(456.9) |
|  | 3-4 | 324 | 1-25 (4.2) | 0-24 (3.2) | 263-16080(2653.2) | 10-2498(248.6) | 58-5840(419.8) |
|  | 3-5 | 253 | 1-16 (4.8) | 0-15 (3.8) | 253-28628(3652.7) | 13-2814(239.2) | 30-9620(490.8) |
| Chr4 | 4-1 | 263 | 1-28 (4.6) | 0-27 (3.6) | 257-23787(3317.3) | 17-3101(226.4) | 65-8707(734.8) |
|  | 4-2 | 261 | 1-20 (5.4) | 0-19 (4.4) | 270-22396(2858.8) | 10-1775(193.8) | 44-3412(395.6) |
|  | 4-3 | 285 | 1-24 (4.8) | 0-23 (3.8) | 217-22049(2853.4) | 12-2603(252.3) | 49-5459(531.9) |
|  | 4-4 | 331 | 1-23 (4.8) | 0-22 (3.8) | 302-17872(2901.4) | 14-4397(239.9) | 27-3615(372.9) |
|  | 4-5 | 503 | 1-26 (6.3) | 0-25 (5.3) | 317-18702(2340.3) | 10-2408(230.9) | 28-6353(308.1) |
| Chr5 | 5-1 | 396 | 1-33 (6.6) | 0-32 (5.6) | 252-14900(2676.8) | 17-3012(213.0) | 53-4007(288.6) |
|  | 5-2 | 392 | 1-22 (5.7) | 0-21 (4.7) | 250-20225(2553.9) | 16-2945(243.6) | 31-5916(290.5) |
|  | 5-3 | 264 | 1-14 (4.5) | 0-13 (3.5) | 291-18950(3077.3) | 9-2855(232.3) | 25-5906(443.7) |
|  | 5-4 | 269 | 1-27 (4.6) | 0-26 (3.6) | 305-21810(2800.6) | 5-2072(227.6) | 35-7228(482.2) |
|  | 5-5 | 300 | 1-23 (4.8) | 0-22 (3.8) | 252-15050(2780.3) | 8-2732(257.2) | 26-2295(323.3) |
| Chr6 | 6-1 | 400 | 1-37 (5.8) | 0-36 (4.8) | 291-20247(2837.5) | 10-2928(219.7) | 29-7129(339.3) |
|  | 6-2 | 315 | 1-27 (4.1) | 0-26 (3.1) | 301-15432(2684.7) | 10-3974(254.1) | 42-8191(365.7) |
|  | 6-3 | 253 | 1-25 (6.7) | 0-24 (5.7) | 288-30412(3385.4) | 16-2350(209.4) | 27-8739(573.2) |
|  | 6-4 | 314 | 1-24 (5.8) | 0-23 (4.8) | 214-29056(2936.7) | 19-2051(219.2) | 27-2259(397.1) |
|  | 6-5 | 423 | 1-22 (5.6) | 0-21 (4.6) | 223-21066(2693.0) | 4-1926(218.4) | 27-2289(318.8) |
| Chr7 | 7-1 | 272 | 1-42 (6.1) | 0-41 (5.1) | 234-18865(3099.5) | 9-2093(191.6) | 27-6395(407.4) |
|  | 7-2 | 301 | 1-26 (5.4) | 0-25 (4.4) | 236-25168(3320.5) | 4-1926(218.9) | 75-5880(501.3) |
|  | 7-3 | 345 | 1-20 (4.7) | 0-19 (3.7) | 300-19469(2766.1) | 13-3038(276.6) | 42-8975(500.4) |
|  | 7-4 | 398 | 1-21 (4.5) | 0-20 (3.5) | 217-15710(2624.5) | 16-2486(244.9) | 50-6733(436.4) |
|  | 7-5 | 420 | 1-40 (6.9) | 0-39 (5.9) | 231-15129(2731.4) | 11-2396(202.8) | 67-8426(333.6) |

Note : The 1-1, 1-2, and so on, in the fragment represents the NO., for example, 1-2 represents the second fragment of chromosome1

### Characterization of genes in different chromosomes of gene density

Gene density was reflected from the two aspects of number and the length of genes. On the one hand, the gene number is the total count of genes contained in the genome of a free-living organism. From more and more genome annotation projects, the estimation of a gene number for each fully sequenced species is becoming increasingly refined. The distribution of genes on different chromosomes based on CDS feature showed that the number of genes on each chromosome was positively correlated to chromosome size, but Chr2 and Chr4 were found to be opposite (Figure2). This phenomenon suggested that the distribution patterns of the genes on various strawberry chromosomes were similar. Totally, the other components besides genes might affect chromosome size in similar modes. The smallest chromosome is Chr1 with 2,736 annotated genes and the largest is Chr6 with 6,453 genes. In addition, the distribution density of genes on Chr1 showed a gradual downward trend from upper to lower end and the genes were highly located in the upper part. The genes in Chr2, 4 and 7 was gradually increased from up to the low end and mainly distributed in the lower part. The genes distribution on Chr3 was a “wavy line” type, which decreased first, then increased and finally decreased. The gene distribution on Chr5, 6 showed a parabolic curve related to slope gradient (Fig.2). As the chart showed that the lengths of the genes could classify the genes into two main groups that were 0–1000bp and 1000–2000bp long, and the 0–1000bp group accounted were more in number. We could see that the length of genes locating at the lower part of Chr1 were relatively longer. Same did those in the upper part of Chr2, 7. The regions with same phenomenon on Chr3, 4, 5, 6 were close to the middle part and did the middle upper part of Chr4. In addition, the distribution of the gene lengths showed some certain regularity: the longer the length of genes in different parts of the same chromosome was the lower the numbers of genes. The lengths of genes in corresponding same part of different chromosomes were approximately same. (Fig.3)

**Figure 2.**
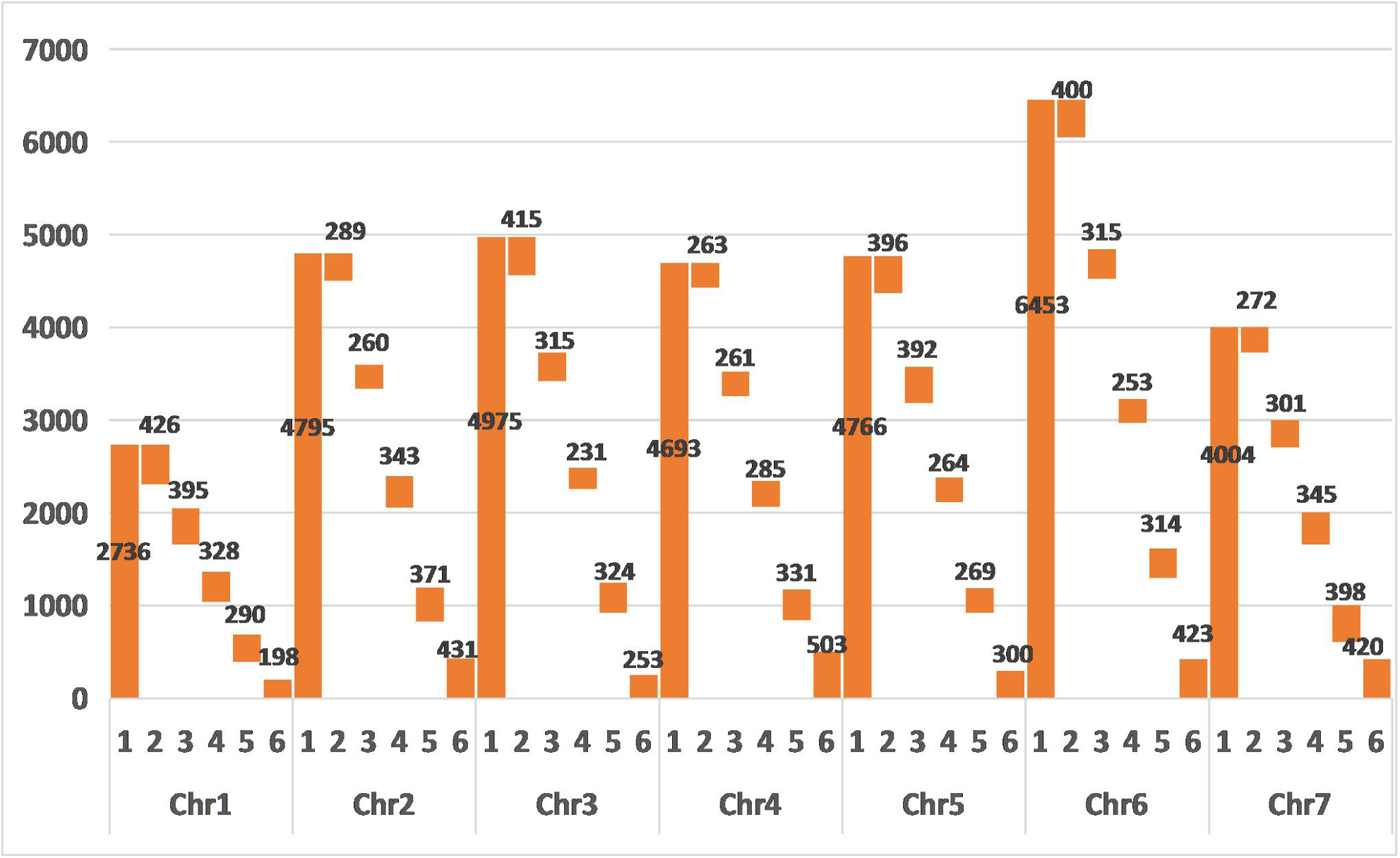
Quantitative distribution of genes on strawberry chromosomes Note: the ordinate of gene number, 1, 2, 3, 4, 5, 6 respectively: total gene number, upper gene number, middle upper gene number, middle gene number, middle lower gene number and lower gene number

**Figure 3.**
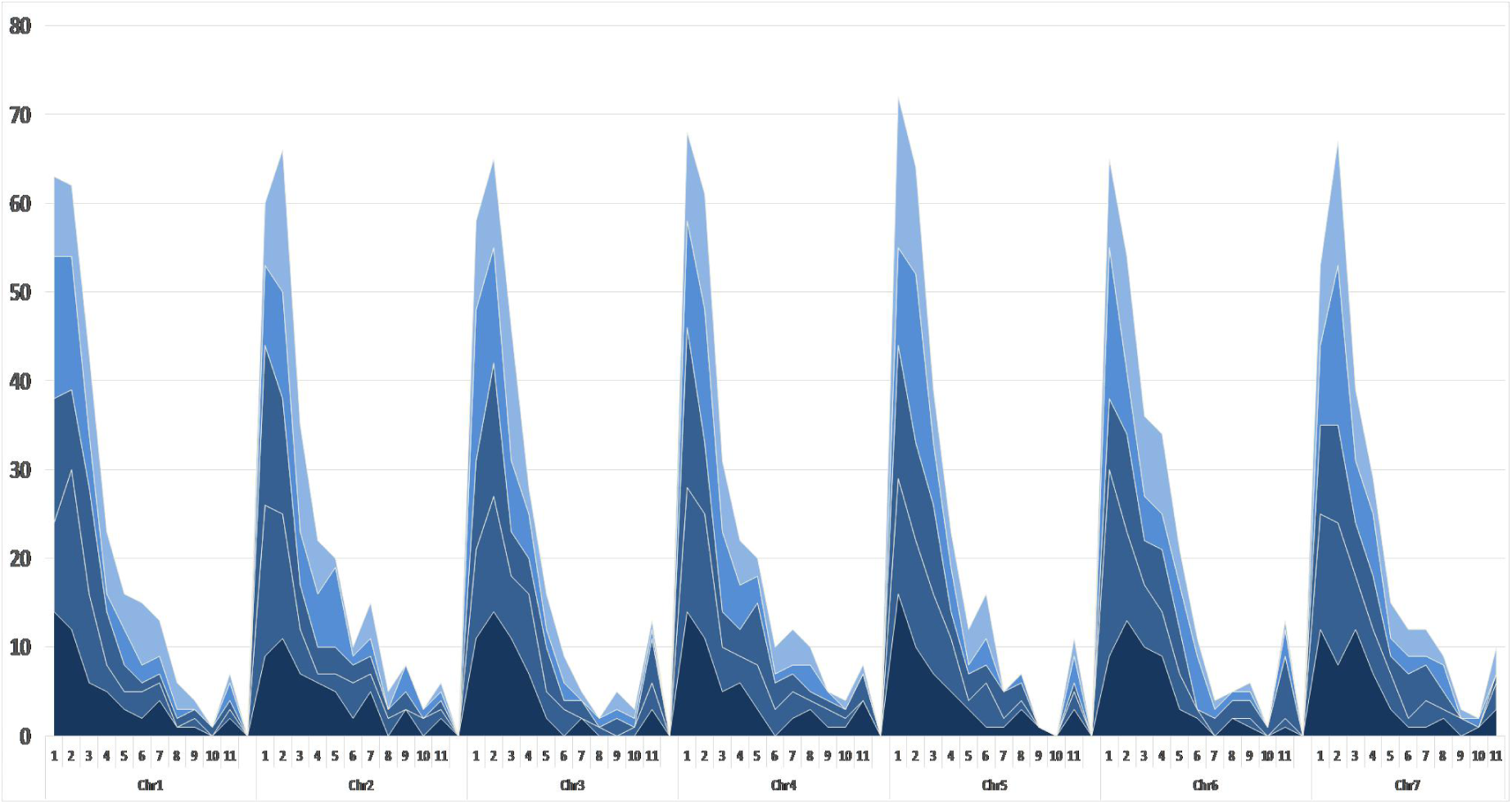
Gene length Note: the abscissa of gene length, 1, 2, 3, 4, 5, 6, 7, 8, 9, 10 respectively:0–1000bp, 1000–2000bp, 2000–3000bp, 3000–4000bp, 4000–5000bp, 5000–6000bp, 6000–7000bp, 7000–8000bp, 8000–9000bp, ≧ 10000bp. Image from top to bottom: upper, middle upper, middle, middle lower and lower

### Characteristic analysis of exons and introns in the selected regions

The description of intron–exon structures is imperative for the theoretical study of the origin and evolution of genes and genomes as well as the function of genes. Therefore, the study of exon and intron structure of strawberry genome is of significance, and the strawberry genome was reinvestigated using the distribution profiles of exon and intron. The analysis of randomly selected 1,750 strawberry genes showed that they contained 9,315 exons and 7,564 introns. The average number of exons in strawberry genes was about 4–6. The features of the exons and introns of the genes locating at all the different chromosome regions were similar. It was found that the change in the number of exons and introns on the chromosome were constant. The length of exons and introns were mainly distributed between a range of 0–200bp, and the change of the same segments of the length of the exons and introns are basically the same. On the other hand, there was increase in the length of exons and introns in a gene,. These results suggest constraints on the splicing machinery to splice out very long or very short introns. It has been reported that the corresponding exon sequences were usually conserved, but the intron sequences were of the opposite situation, and the intron evolution was much faster than that of the exons, since the intron does not have to produce a protein with a useful sequence pressure [19–21]. In our study the similarity of exon lengths was much higher than that of introns on each chromosome (Table 1, Fig. 4).

**Figur 4.**
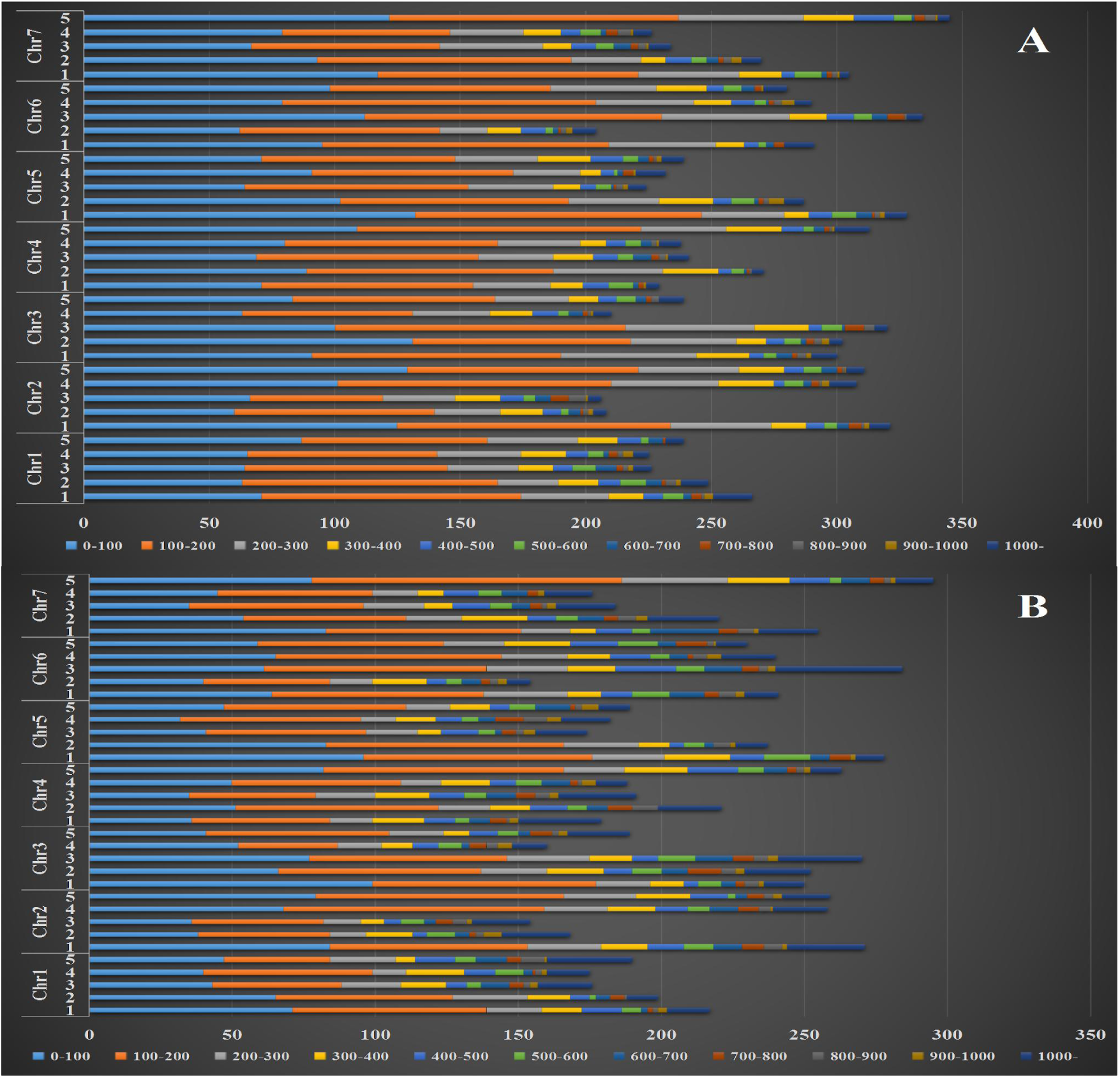
The length of exons (A) and introns (B) Note. The abscissa of intron number, 0–100, 100–200, 200–300, 300–400, 400–500, 500–600, 600–700, 700–800, 800–900, 900–1000, 1000-respectively the length of introns (in bp). the ordinate of chromosome, 1,2,3,4,5 respectively: upper, middle upper, middle, middle lower and lower

It was observed that the average length of exon was about 229 bp, and the length of exons in all regions ranged from 4bp to 4,397bp. About 82% of the exons on each chromosome were lower than 300 bp in length. Fewer than 4% of the exons were over 1,000 bp in length. The lengths of introns were longer than those of exons, and the average size is about 413 bp, with a range from 24bp to 9,757bp. About 8.7% of introns were more longer than 1,000 bp. However, most (66%) of the introns on each chromosome were shorter than 300 bp. As shown in Figure 4 about the distributions of the lengths of exons and introns in the different regions selected on the chromosome. For Chr1 and 6, the longest of exons was found in the first and second segments. In Chr2, 4 and 7 the longest of exons was detected in the third and fourth paragraphs. Chr3 was mainly in the fourth and fifth segment of the longest of exons, Chr5 was the fifth paragraph. In addition, the introns of the genes in the fifth segment on Chr1 and 3 were the longest, and the situation of Chr2 and 7 was their second and third paragraphs had the genes with longest introns. Same did the first segment on Chr4, and the third and fourth segments on Chr5 and 6. These results demonstrated that there was some negative correlation between the lengths of the exons and the introns (Table 1, Fig. 4).

### Distribution of intronless gene

In this study, 6,500 (20.05%) intronless genes were also identified. The numbers of intronless genes on the 7 chromosomes ranged from 669 to 1,354. The ratios of intronless genes to the total genes on Chr1, 2, 3, 4, 5 were roughly constant about (12.23%). The ratios on Chr6, 7 were higher, and they were 17.71% and 20.83%. In order to test whether the number of intronless genes on each chromosome were correlated with the gene numbers and the chromosome lengths in strawberry, linear equation was established, and the result showed that the number of intronless genes were nonlinear with the number of genes on each chromosome (R^2^ =0.1303; Fig. 5A) as well as with the corresponding chromosome lengths (R^2^ = 0.0057; Fig. 5B). It was also found that the distributions of intronless genes in different regions even on same chromosome were different. For example, Chr1 owned a large percentage of intronless genes (27.27%), with reducing from the 1 to the 5. In Chr2, the numbers of intronless genes were evenly distributed. The distribution of intronless number on Chr3, 5 and 6 showed a trend of parabola, and in the three segments of the number genes recorded the least. The numbers of intronless genes were increasing from top to bottom on Chr4. Also, the distribution of intronless number on Chr7 showed a similar trend of parabola, further, we noticed a more number of genes present in the three segments (Fig. 5C).

**Figur5.**
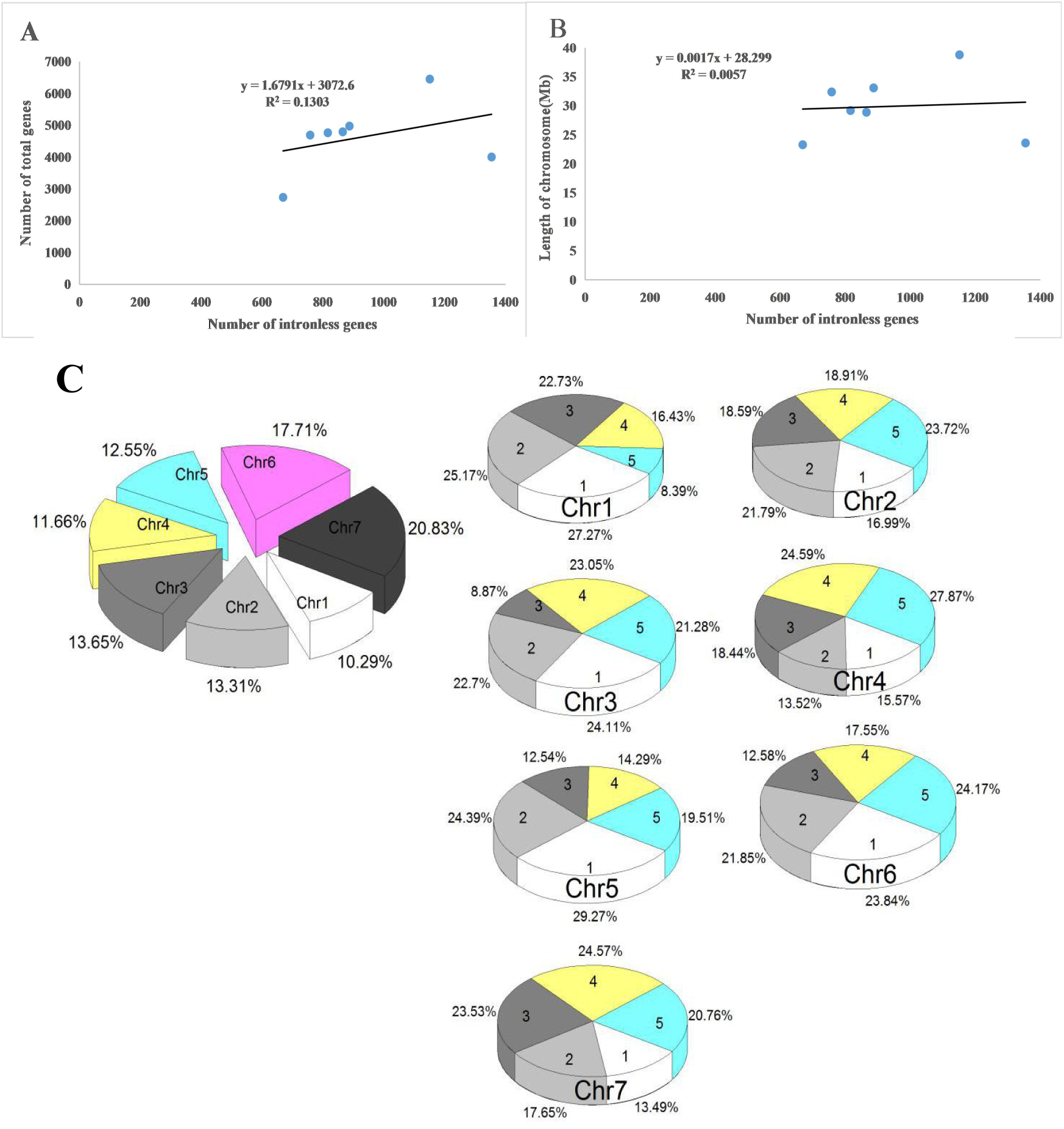
Linear correlation among intronless genes, total genes, and the length of chromosome. (A) The linear regression equation between the number of intronless genes on each chromosome and the number of genes (including intronless and intron-containing genes) on the chromosome. (B) The linear regression equation between the number of intronless genes on each chromosome and the length of their respective chromosome.(C) Distribution of intronless genes.1,2,3,4,5 respectively: upper, middle upper, middle, middle lower and lower

### Expression analysis of the genes

The expression levels of genes on different chromosomes were different, the genes in different segments of the same chromosomes were also different. However, the expression levels of the genes in similar segments of the different organization on the chromosomes were consistent. As shown in the figure, in Chr1 the expression levels of genes in the upper and middle upper parts showed high expression, while in middle lower and lower parts the expression was low. Expression of the genes in middle upper, lower parts of the higher and other parts were lower on Chr2. For Chr3, the higher expression level on the upper, middle and middle lower parts, whereas in the middle upper and lower part it was lower. In Chr4, the middle upper and middle lower part of the gene expression is higher in comparison to other parts which recorded lower expression. Chr5 was mainly in the middle and middle lower part of the highest expression and other parts recorded a lower expression. The lowest of gene expression was noticed in the middle part and other parts of the Chr6 showed high expression. In Chr7, the expression of the upper and lower genes were higher and it was relatively low in other parts (Fig.6)

**Figure 6.**
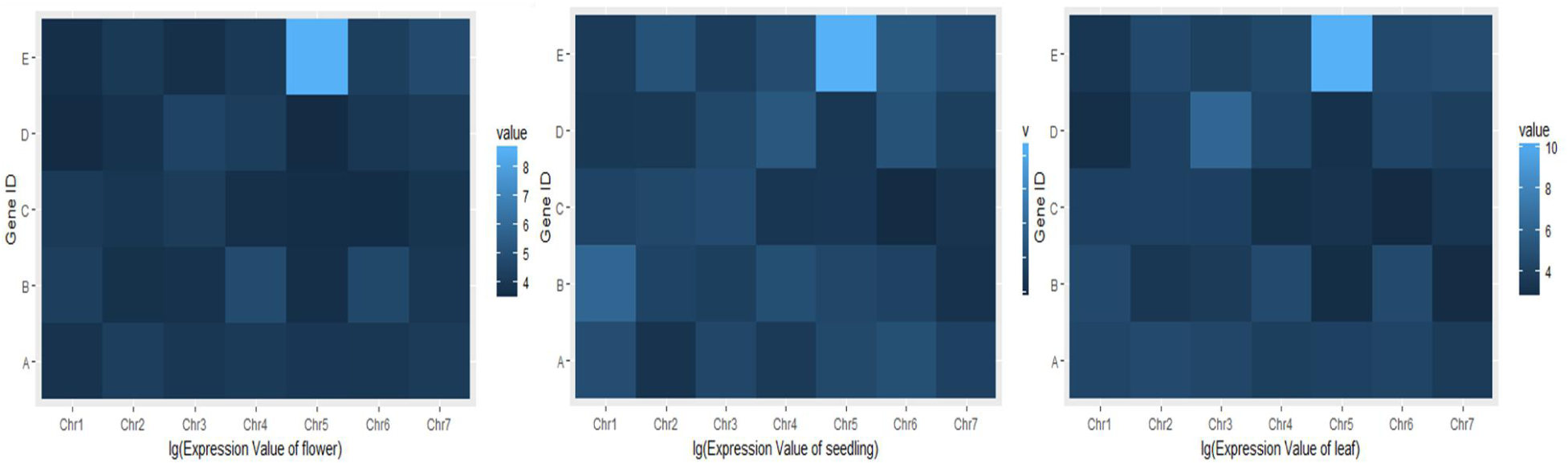
Different parts of the gene expression Note: A,B,C,D,E,respectively: upper, middle upper, middle, middle lower and lower

### Correlation analysis between each index

In the present study, there was a negative correlation observed between the number of genes and gene length, suggesting that when the gene numbers on a chromosome was higher the lengths of genes and the gene density will be shorter, also we found that there was a weak positive correlation between the number of genes and the number of exons and introns, while a good positive correlation relationship was noticed between the number of genes and intronless genes, the more the number of genes where intronless gene are also more. In addition, intronless gene is also affected by other factors, but the number of exons and introns has little effect on it. For the number of exons on the gene and its length we can see that they are negatively correlated, and the number and length of introns is consistent with the exon. However, affect the number of introns on length is less than the exons, which indicates that the length of the intron also affects the other important factors. Finally, the length of the intron is important for the length of the gene, and the length of the gene depends primarily on the length of the intron and does not vary much from the length of the exon. This means that very long genes are due to their long introns, rather than the need to encode longer products (Fig.7)

**Figure 7.**
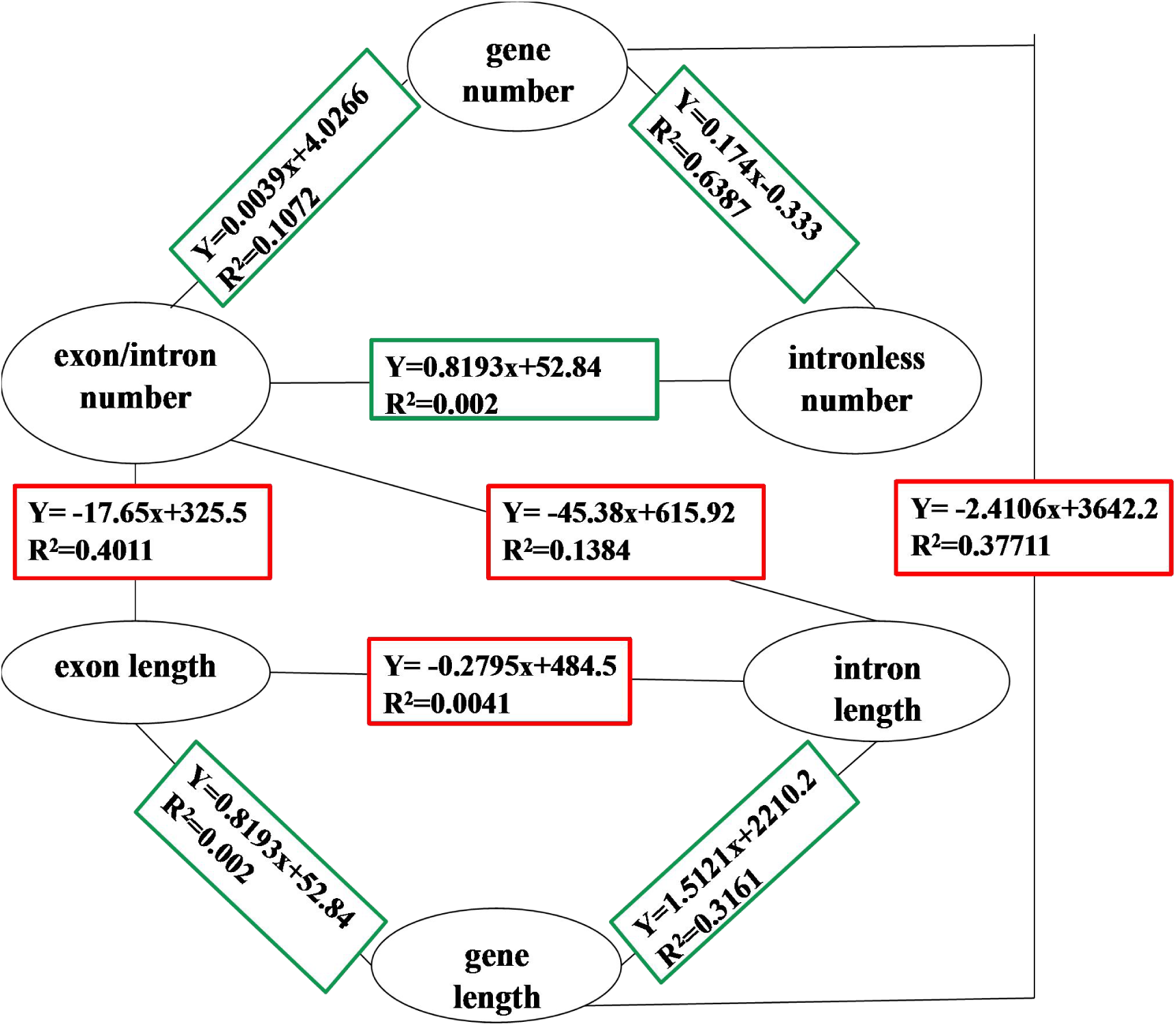
Correlation analysis between each index Note: Positive and negative correlations are indicated by green and red lines, respectively. The numbers above and below each line indicate the correlation coefficient formula and R2 of the significance, respectively.

### Prediction of centromeric position of strawberry chromosomes

The centromere is a chromosomal constriction region that contains meiosis or mitotic spindle binding sites. Mapping centromere locations in plant species provides essential information for the analysis of genetic structures, genetics and population dynamics. The location of centromere has been an interesting and important work. In the last two decades, intensive studies on the biology of centromeric regions in plants are being conducted, and considerable success has been achieved [22–27]. The structures of chromosomes are characteristic to the position of centromere on each chromosome exhibiting lower number of intronless genes and low gene density [28–29], the level of a genes expression and the length of its introns were correlation and reported [17]. Also, the lengths of genes were examined in a set of constitutively expressed genes [18]. Based on the characteristic features of chromosome, this study used the intron length, intronless genes, gene length and gene expression to predict the position of the strawberry centromere (Table 2).

**Table 2.**
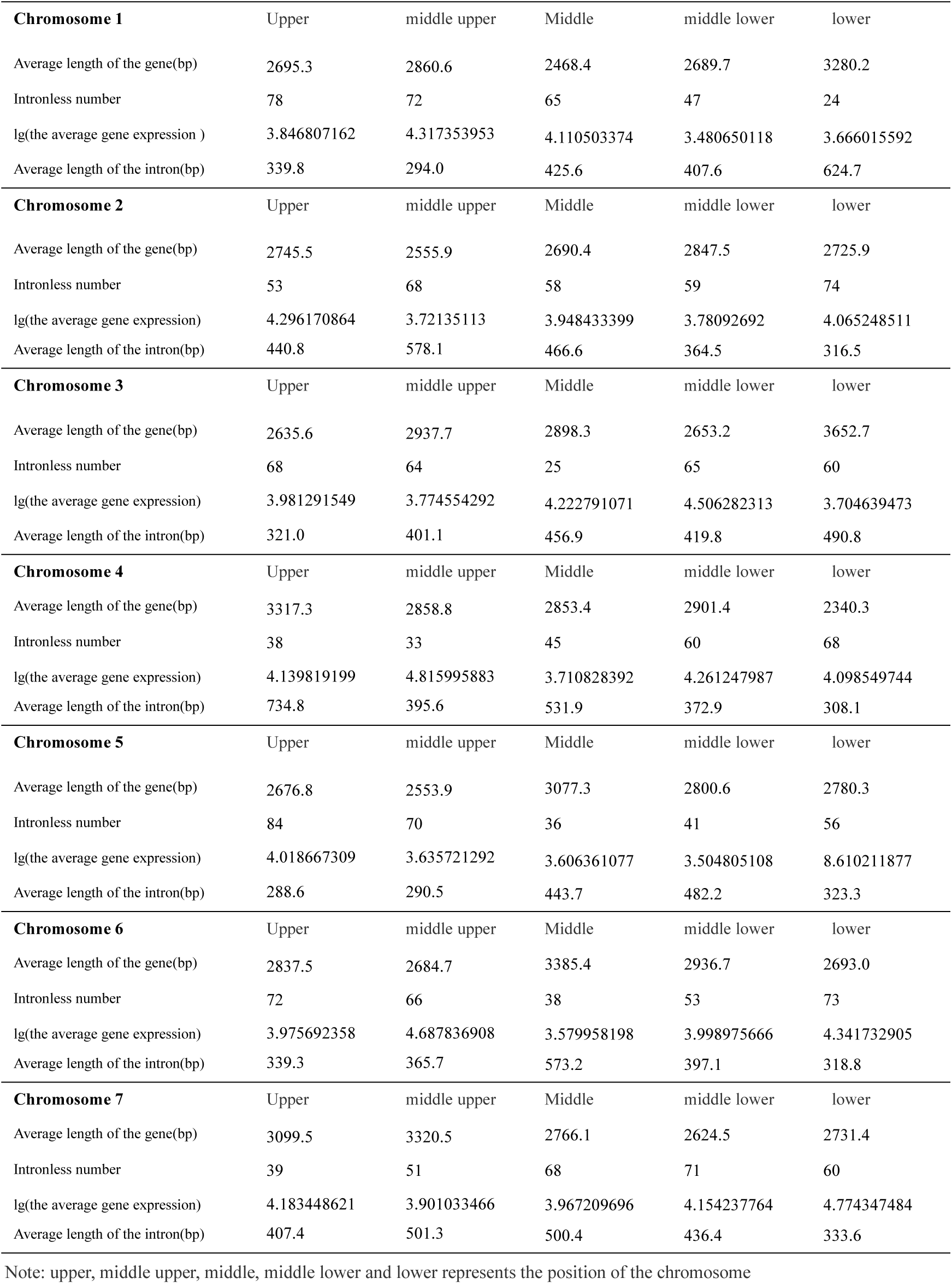
Strawberry centromere position-related data

The prediction analysis gave the rough position of the centromere of the strawberry seven chromosomes (Figure 8). From the figure we predict that Chr1 and 7 are telocentric chromosomes, Chr2 and 3 is acrocentric chromosome. Chr4 are submetacentric chromosomes, Chr5 and 6 are metacentric chromosomes. Therefore, we have predicted the centromere position of strawberries in combination with previous studies, and these findings will also help to optimize breeding strategies based on the sexual hybridization and also contribute to a better understanding of strawberry genetics.

**Figure 8.**
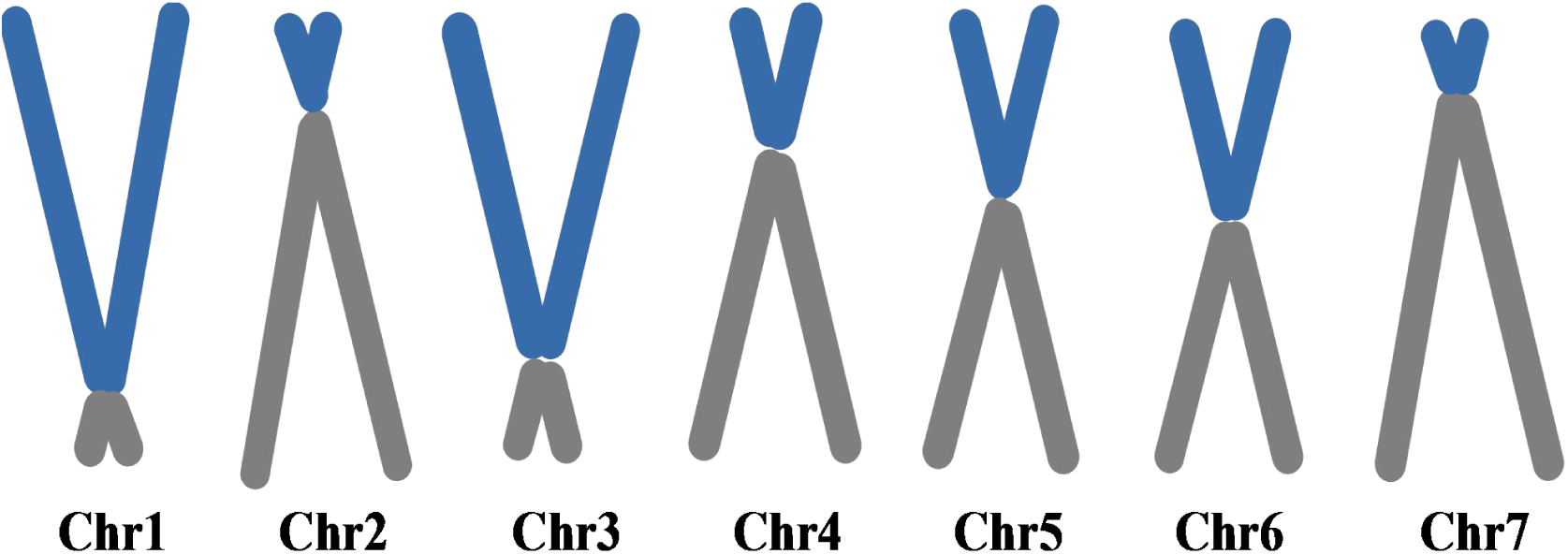
According to the genetic density and other structural features of the relative position of centromere

## Discussion

The rapid development of bioinformatics and genomics makes the analysis of genome sequence characteristics a hot spot. There are many analyzes of genomic sequence characteristics, such as Chiaromonte et al. they reported the effects of gene length and proximity to genome-wide expression levels [18]; Roy et al. studied the effect of intron length on exon creation ratios during the evolution of mammalian genomes [30]; The chromatin organization marks exonintron structure was documented by Schwartz et al. [31]. Strawberry lacks substantial genomics-level resources [32], for the rapid development of genetic information but now does not match. We know the strawberry is small stature, rapid seed-to-seed cycle, transformability and miniscule basic genome make strawberry an attractive system to study processes related to plant physiology, development and crop production. Therefore, in this study we explored the sequence characteristics of different sections of strawberry (*Fragaria vesca*) genome. This work will pave a a new era of applying genetic information to a deeper understanding of strawberries and used to enhance the agricultural production. Making genetic information and genomics applied in many aspects of strawberry production, which means that the era of application of genetic information in crop production has arrived.

In this study, about 32,422 genes on strawberry chromosomes were used as experimental material and analyzed by various bioinformatics software to select different regions on chromosome to analyze and predict the position of strawberry centromere. Here we choose 2M five sections of the region to facilitate the analysis, each section are randomly selected and the results found that the characteristics of the five sections can also explain the problem. Our results show that exon length are distributed much more tightly than introns, this implies that exons lengths are limited to a shorter range than introns length. These results are consistent with previous observations for eukaryotic genomes [15]. In addition, the length of the gene was positively correlated with the length of the exon and intron. The length of the gene depends mainly on the length of the intron, and the length of the exon is much altered. Very long genes are due to their introns, rather than the need to encode longer products. This study with Swinburne and Miguez demonstrated that introns contribute significant length to genes, break the protein coding information so that it can be alternatively processed into different splice isoforms introducing a unique level of regulation [33]. It is worth mentioning that according to previous studies, we know that the position of maize centromere on each chromosome exhibit lower number of intronless genes [28]. In addition, Pablo Aleza et al. reported that citrus centromere has a low gene density [29]. In this paper, we also use the lower gene expression as a centromere position prediction. A strong correspondence was reported between the level of a gene’s expression and the length of its introns [17]. In another work, gene length was examined in a set of constitutively expressed genes [18]. Therefore, we have positively predicted the centromere position of strawberry. These findings will also help to optimize breeding strategies based on sexual hybridization and also contribute to a better understanding of strawberry genetics. The position of centromere on strawberry might be affected by many factors, our forecast also only through existing research on other crops. So, further research is needed in this line of work.

In summary, the study of strawberry genome characteristics could help to understand the evolutionary patterns of related genes and genomes. Genomic and bioinformatics validation of the wild strawberry genome characteristics, for the future more in-depth understanding of the strawberry genome lay a solid foundation. It is expected that the results reported herein will be of particular value to plant biologists interested in the studied genes and the traits to which they may relate, and to molecular breeders pursuing the genetic improvement of strawberry and/or other members of the Rosaceae family.

## Conclusions

About 32,422 genes on strawberry chromosomes were evaluated, and a number of bioinformatic softwares were used to analyze the characteristics of genes, exons and introns, expression of genes in different regions on the chromosomes. Also, the positions of strawberry centromeres were predicted. The results of this investigation could definitely provide a significant foundation for further research on function analysis of gene family in Strawberry.

## List of abbreviations

Chr1: chromosome 1
Chr2: chromosome 2
Chr3: chromosome 3
Chr4: chromosome 4
Chr5: chromosome 5
Chr6: chromosome 6
Chr7: chromosome 7

## Declarations

### Ethics approval and consent to participate

Not applicable

Consent for publication

Not applicable

Availability of data and material

Not applicable

### Authors’ contributions

FJG, SGLF conceived and designed the experiments. LA performed the data analysis and wrote the manuscript. FJG and Sudisha Jogaiah revised the manuscript. All authors reviewed and approved the manuscript.

### Acknowledgements

The authors wish to thank Nanjing Agricultural University, Jiangsu, China. Chen Lide, Liu Zhongjie, Cui Mengjie, Shangguan Lingfei, Jia Haifeng and Fang Jinggui.

## Additional files

**Additional file 1: Figure S3.** Gene length (XLS 34 kb)

**Additional file 2: Figure S4.** Exon length (XLS 31 kb)

**Additional file 3: Figure S4.** Intron length (XLS 31 kb)

**Additional file 4: Figure S5.** Intronless number (XLS 32 kb)

**Additional file 5: Figure S6.** Different parts of the gene expression(XLS 22 kb)

